# Mosaic and cocktail capsid-virus-like particle vaccines for induction of antibodies against the EPCR-binding CIDRα1 domain of PfEMP1

**DOI:** 10.1101/2024.04.02.587801

**Authors:** Ilary Riedmiller, Cyrielle Fougeroux, Rasmus W. Jensen, Ikhlaq H. Kana, Adam F. Sander, Thor G. Theander, Thomas Lavstsen, Louise Turner

## Abstract

The sequestration of *Plasmodium falciparum-*infected erythrocytes to the host endothelium is central to the pathogenesis of malaria. The sequestration is mediated by the parasités diverse *Plasmodium falciparum* erythrocyte membrane protein 1 (PfEMP1) variants, which bind select human receptors on the endothelium. Severe malaria is associated with PfEMP1 binding human endothelial protein C receptor (EPCR) via their CIDRα1 domains. Antibodies binding and inhibiting across the sequence diverse CIDRα1 domains are likely important in acquired immunity against severe malaria. In this study, we explored if immunization with AP205 bacteriophage capsid-virus-like particles (cVLPs) presenting a mosaic of diverse CIDRα1 protein variants would stimulate broadly reactive and inhibitory antibody responses in mice. Three different mosaic cVLP vaccines each composed of five CIDRα1 protein variants with varying degrees of sequence conservation of residues at and near the EPCR binding site, were tested. All mosaic cVLP vaccines induced functional antibodies comparable to those induced by matched cocktails of cVLPs decorated with the single CIDRα1 variant. No broadly reactive responses were observed. However, the vaccines did induce some cross-reactivity and inhibition within the CIDRα1 subclasses included in the vaccines, demonstrating potential use of the cVLP vaccine platform for the design of multivalent vaccines.

## Introduction

Despite concerted efforts to control malaria, the disease remains a significant public health challenge, with severe cases posing a significant threat, especially to children [1]. In regions with high malaria transmission, immunity to severe malaria develops early in life, after a limited number of malaria episodes [2]. Central to malaria pathogenesis is the tissue sequestration of parasite-infected erythrocytes, mediated by the *Plasmodium falciparum* erythrocyte membrane protein 1 (PfEMP1) [3]. Of particular significance for severe outcomes of malaria is the interaction between the CIDRα1-bearing PfEMP1 and the endothelial protein C receptor (EPCR) [4]. The selective pressure exerted by the host immune system has led to the emergence of PfEMP1 variants exhibiting substantial diversification in their amino acid sequences. Nonetheless, naturally acquired antibodies have demonstrated the capacity to overcome this obstacle [5], either through the development of a limited number of antibodies that cross-react with variants within the sequence-defined subgroups of CIDRα1 domains (CIDRα1.1-1.8) or broadly reactive antibodies that recognize all CIDRα1 variants. Such broadly reactive and inhibitory antibodies may target conserved epitopes on CIDRα1 domains, arising from the biochemical and structural constraints to retain EPCR binding.

Prior studies on the immunogenicity of CIDRα1 domains have included recombinant CIDRα1 protein domains as single variants or cocktails of multiple variants, live attenuated adenoviruses encoding CIDRα1 or capsid-virus-like particles (cVLPs) decorated with a single CIDRα1 protein variant [6–8]. These strategies have consistently demonstrated the elicitation of anti-CIDRα1 antibodies capable of inhibiting EPCR binding, not only to the CIDRα1 used in the vaccine formulation but also, to a lesser extent, towards other CIDRα1 variants belonging to the same CIDRα1 sequence subgroup.

In this study, we investigated if broadly inhibitory CIDRα1 antibodies could be induced through immunization with cVLPs decorated with five different CIDRα1 variants. These were predicted to share an epitope due to a common structural fold presenting residues critical for EPCR binding. The residues outside the common epitope differed among the recombinant domains coating the cVLPs. This “mosaic cVLP” strategy rested on the hypothesis that the ordered and repetitive display of multiple proteins on a single cVLP can promote the stimulation of B-cells expressing cross-reactive and inhibitory antibodies. This would take place through the direct activation of B-cells by cross-linking their receptors, which recognize the shared epitope of adjacent heterotypic CIDRα1 [9,10].

We find that mosaic cVLPs decorated with five sequence-diverse CIDRα1 variants, selected from the natural pool of CIDRα1 sequences to share a few surface-exposed residues at pre-defined positions, elicited functional antibodies with a breadth comparable to immunization with a cocktail of different homotypic cVLPs. While we find no indication of induction of broadly reactive responses, the induced cross-reactivity within CIDRα1 subclasses, indicates that a multivalent vaccine, in principle, is feasible.

## Results

### Selection of CIDRα1 variants for cVLP vaccine design

The EPCR-binding CIDRα1 domains group into subsets CIDRα1.1, 1.4-1.8 [5,11]. The six subgroups likely reflect an antigenic diversification of the protein family resulting from antigenic drift and a limited recombination occurrence within the gene elements encoding the CIDRα1 domain [12]. However, the structural and sequence diversity of the EPCR-binding site of CIDRα1 domains are restricted to maintain high affinity for the receptor [5]. Here we attempted to exploit this by designing three mosaic cVLPs, which differed in the conservation of surface-exposed amino acids at and near the EPCR-binding site. Specifically, CIDRα1 variants were selected based on their amino acid conservation at the seven positions on the EPCR-binding helix (EB helix) mediating direct interaction with EPCR, and at six positions on the adjacent EPCR binding supporting (EBS) helix (Figs 1A and B). CIDRα1 variants were selected from a sequence database of ∼3000 *P. falciparum* genomes, containing 21 764 different naturally occurring CIDRα1 sequences [13]. Within this dataset, 312 unique sequences were identified across the 13 specified positions (Fig 1C).

**Fig 1.**
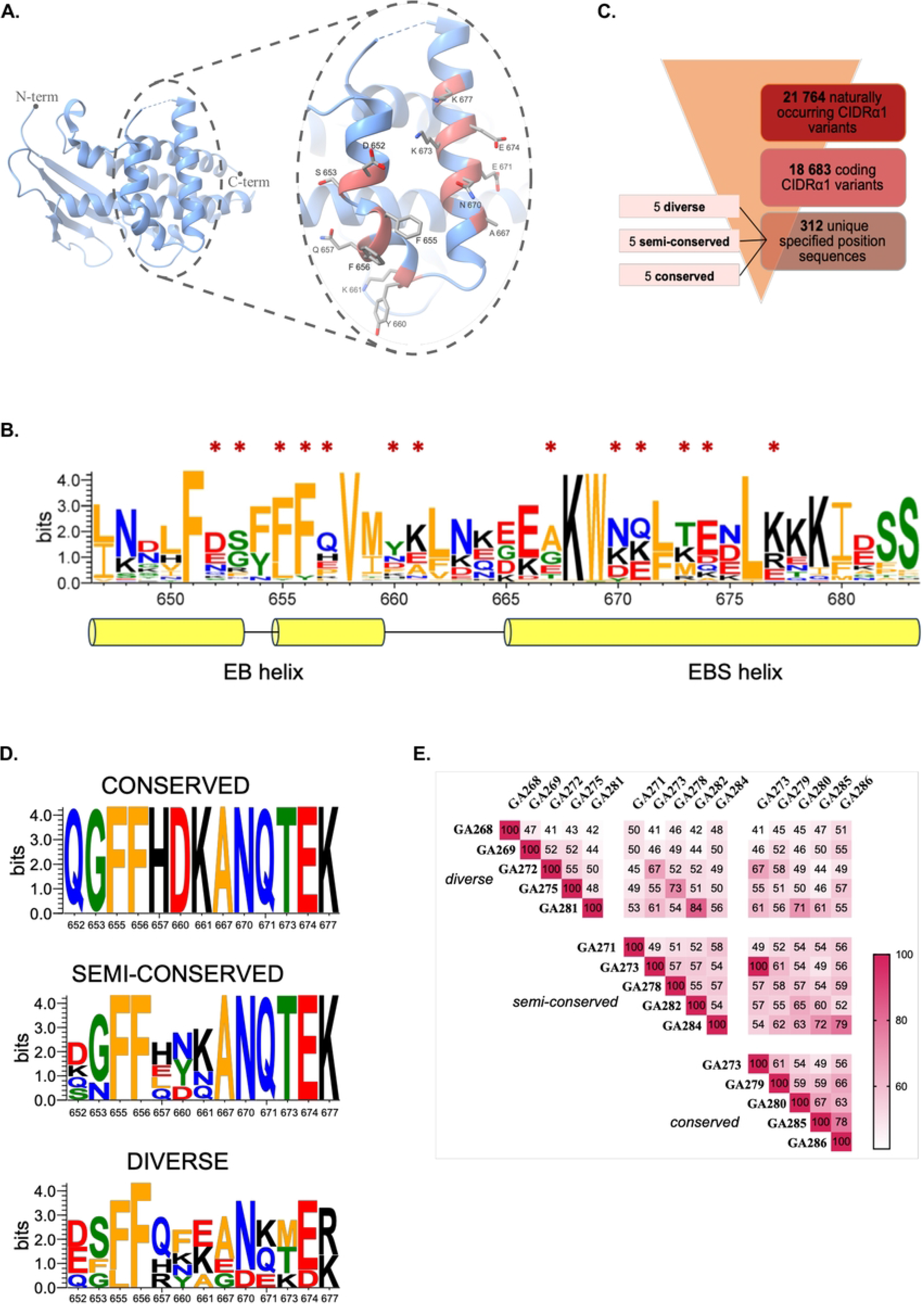
Selection of naturally occurring CIDRα1 sequences for the design of CIDRα1-cVLP vaccines. **A**. Structure of the HB3VAR03 CIDRα1.4 domain with close-up on the EPCR binding (EB) and EPCR binding supporting (EBS) helices [5]. Highlighted in red are the amino acids kept constant among CIDRα1 variants chosen for the *mosaic-conserved-*cVLP vaccine. **B**. Sequence conservation LOGO of 963 naturally occurring CIDRα1 sequences. Asterisks indicate the positions kept constant among CIDRα1 variants chosen for the *mosaic-conserved-*cVLP vaccine. Marked residues on the EB helix also interact directly with EPCR. **C**. Schematic representation of the selection process resulting in the identification of the CIDRα1 variants used in this study. **D.** Sequence LOGO of the 13 selected amino acid positions among the CIDRα1 variants selected for the three mosaic cVLP vaccines. **E.** Pairwise sequence identity of the 14 CIDRα1 variants selected for the mosaic cVLP vaccines. One variant (GA273) is shared between the *conserved* and *semi-conserved* cVLP vaccines.

For the most conserved mosaic cVLP vaccine, groups of CIDRα1 sequences sharing amino acids at all 13 positions (Fig 1D) were searched among 21 764 naturally occurring CIDRα1 variants to identify five, which differed the most in the surrounding amino acid composition. The variants chosen represented subgroups CIDRα1.4/5/7 and had an average pairwise sequence identity of 61%. For the most diversified mosaic cVLP, five randomly chosen CIDRα1 variants representing five of the six CIDRα1 subgroups (all but CIDRα1.8) were chosen (Fig 1D). These variants had an average pairwise sequence identity of 47%. In addition, a semi-conserved mosaic cVLP, was designed with five variants representing five different CIDRα1 subgroups (CIDRα1.4-8) and sharing amino acids only at the seven positions on the EBS helix (Fig 1D). This group of proteins shared on average 55% amino acids. In total, the three mosaic cVLP vaccines were composed of 14 different CIDRα1 sequence variants, with varying degrees of pairwise sequence identity (Figs 1E and S1).

### Formulation and characterization of cVLP vaccines

Recombinant CIDRα1 proteins were produced in insect cells fused with a SpyCatcher (SpyC) domain to the N-terminus, as this design has previously produced stable EPCR-binding proteins capable of eliciting inhibitory antibodies [6]. The SpyC domain enables direct coupling of the proteins to pre-assembled *Acinetobacter phage* 205 (AP205) cVLPs with SpyTag peptides genetically fused to the N-terminal of the capsid protein [14,15]. The recombinant SpyC-CIDRα1 proteins were coupled to SpyT fused AP205 cVLPs individually and together to generate homotypic and mosaic cVLPs, respectively, as specified in Fig 2A. All mosaic and most homotypic formulations showed stable conjugated cVLPs (Figs 2B and S2). Three unstable homotypic cVLP formulations (cVLP-GA275, -GA279 and -GA284) were excluded from the study. Dynamic light scattering analysis was conducted to further examine vaccine particle sizes and propensity to aggregation. The analysis confirmed successful coupling albeit with a relatively high polydispersity (20-81% Pd) indicating a propensity for aggregation in all cases (Figs 2C and S3).

**Fig 2.**
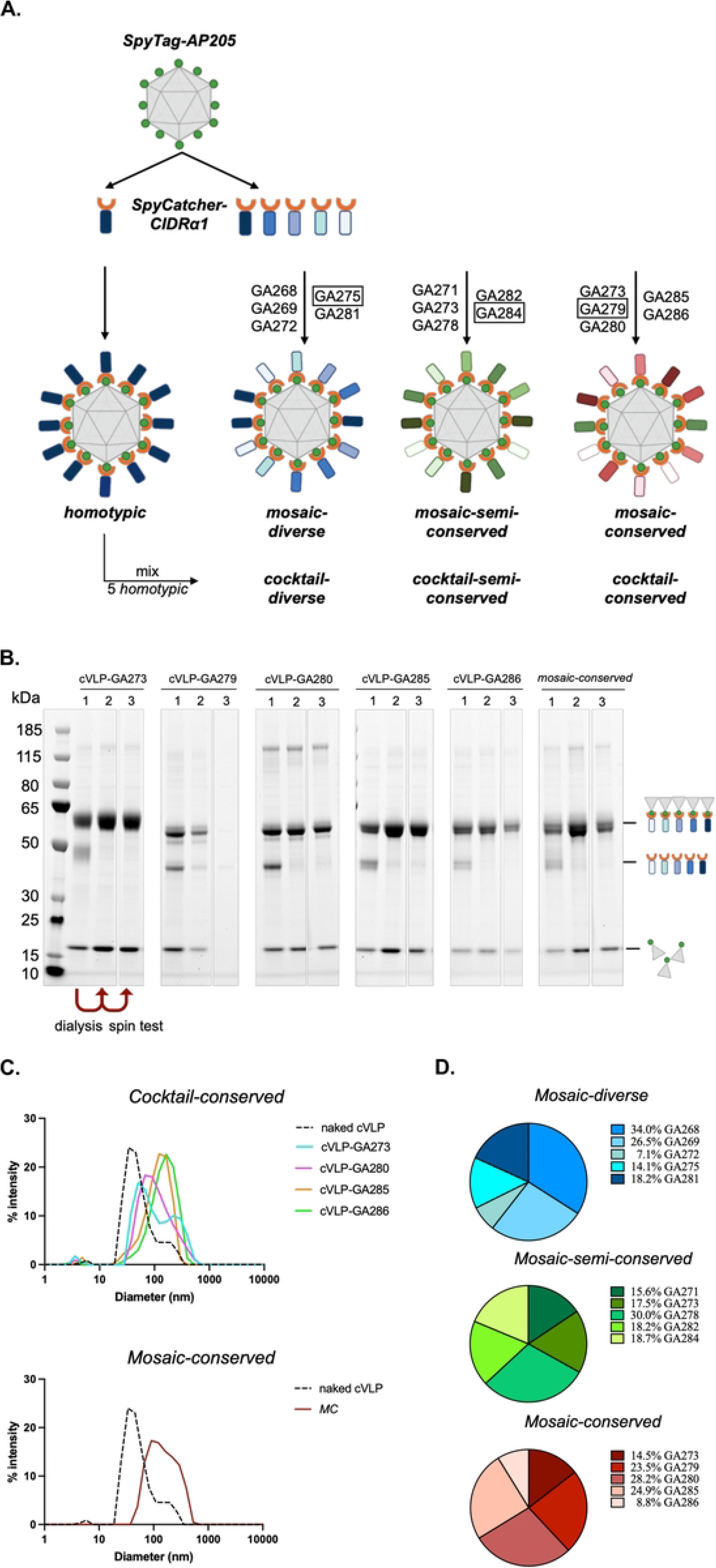
Formulation and characterization of vaccines. **A.** Schematic representation of vaccine formulations. Coupling of individual CIDRα1 proteins on cVLPs gives rise to homotypic particles, which are mixed to obtain three different cVLP cocktails (*cocktail-diverse*, *cocktail-semi-conserved, cocktail-conserved*) matching composition of the mosaic cVLPs. Coupling of five variants on the same cVLP results in mosaic cVLP vaccines (*mosaic-diverse*, *mosaic-semi-conserved, mosaic-conserved*). Coupling of homotypic cVLPs with CIDRα1 variants highlighted by rectangles was unsuccessful thus, the equivalent homotypic cVLPs lack from the corresponding cocktails. **B**. SDS-PAGE of five homotypic cVLPs and *mosaic-conserved*-cVLPs. First lane (1) represents the samples pre-dialysis. The bands correspond to Tag-cVLP (16.5 kDa), unbound Catcher-CIDRα1 (∼40 kDa) and coupled CIDRα1-cVLP (∼56 kDa). The second lane (2) represents the samples post-dialysis and before spin test. The third lane (3) represents the samples post-dialysis and post-spin test. SDS-PAGE of the remaining samples is reported in S2 Fig. **C**. Dynamic-Light-Scattering analysis of *cocktail-conserved*-cVLPs and *mosaic-conserved-*cVLPs. Similar data for the remaining cVLP vaccines is reported in S3 Fig. Naked SpyT-cVLPs (dashed line) predominant population shows 65.7 nm with 81.2% Pd. *Cocktail-conserved-*cVLPs indicate a population size of 69.3-162.9 nm with 38.9-72.4% Pd and *mosaic-conserved*-cVLPs show 170.6 nm and 58.7% Pd. **D**. Distribution of CIDRα1domains coupled to cVLPs as indicated by mass spectrometry analysis (details are reported in *supplementary methods)*. Abbreviations: MC, mosaic-conserved.

For homotypic cVLPs, coupling efficiencies were estimated at ∼41-62% (S4 Fig), translating to ∼73-111 recombinant CIDRα1 proteins conjugated to each cVLP, in accordance with previous reports [16,17]. Bottom-up, label-free Tandem Mass Spectrometry was used to assess coupling efficiency of different protein variants on the same mosaic cVLP. This confirmed successful conjugation of the five different CIDRα1 domains on each of the three mosaic cVLPs and returned an estimate of their relative distribution on the cVLPs (Figs 2D and *supplementary methods*). While these data do not formally confirm the presence of all five proteins on the same cVLP, their even distribution in the total formulation is suggestive of mosaics formation. Among the three, the *mosaic-semi-conserved*-cVLPs showed the most even CIDRα1 distribution, while analysis of the *mosaic-diverse*-cVLPs and *mosaic-conserved*-cVLPs indicated a 3-4 times difference in the abundance of the most and least prevalent CIDRα1 variant, probably reflecting small alterations in coupling efficiencies.

### Antigen binding and inhibition capacity of cVLP vaccines induced antibodies

Homotypic cVLPs were mixed to generate cocktails matching the CIDRα1 variant composition of each of the three mosaic cVLPs, except that each cocktail lacked one variant (cVLP-GA275, -GA284 and -GA279) due to unsuccessful coupling to cVLPs (Fig 2A). Groups of 10 BALB/c mice (only five mice for *cocktail-diverse-*cVLPs) were immunized twice with 3-weeks interval with cocktail-and mosaic cVLP vaccines without adjuvant. In addition, groups of five mice were immunized in a similar scheme with cocktails of recombinant CIDRα1 proteins matching the mosaic cVLP compositions and six of the single homotypic cVLPs (Table 1). IgG was purified from pooled sera of mice from each immunization group, collected two weeks post booster dose. Reactivity and EPCR-binding inhibition ability of purified IgG was assessed towards a panel of 25 CIDRα1 domains, in a bead-based multiplexed assay [6].

**Table 1.**
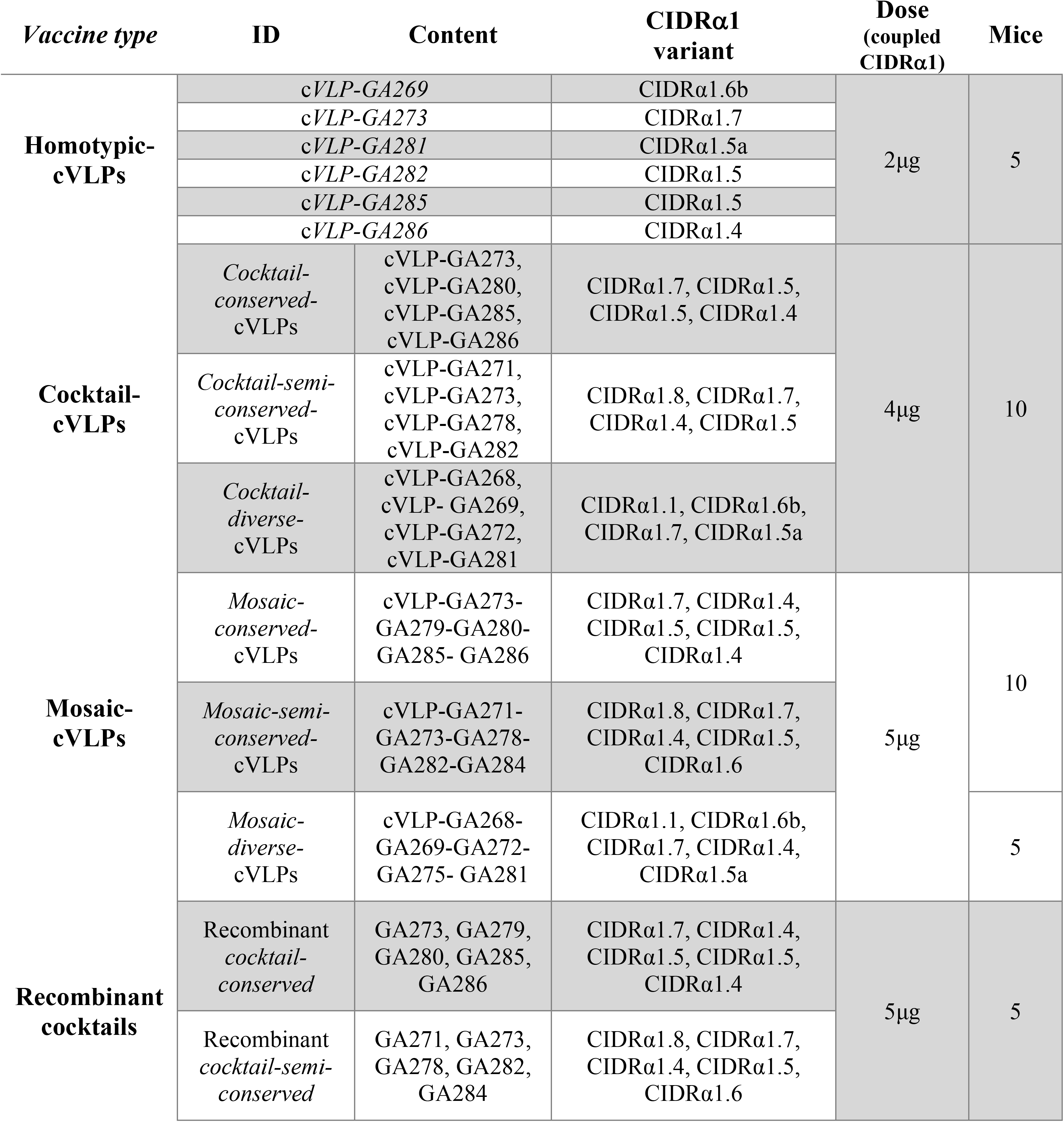
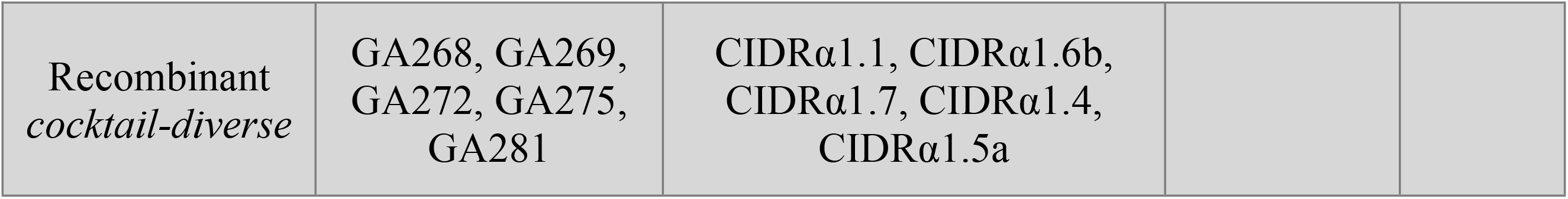
Vaccination scheme.

| *Vaccine type* | **ID** | **Content** | **CIDRα1 variant** | **Dose (coupled CIDRα1)** | **Mice** |
| --- | --- | --- | --- | --- | --- |
| **Homotypic-cVLPs** | *cVLP-GA269* | | CIDRα1.6b | 2μg | 5 |
| | *cVLP-GA273* | | CIDRα1.7 | | |
| | *cVLP-GA281* | | CIDRα1.5a | | |
| | *cVLP-GA282* | | CIDRα1.5 | | |
| | *cVLP-GA285* | | CIDRα1.5 | | |
| | *cVLP-GA286* | | CIDRα1.4 | | |
| **Cocktail-cVLPs** | *Cocktail-conserved-cVLPs* | cVLP-GA273, cVLP-GA280, cVLP-GA285, cVLP-GA286 | CIDRα1.7, CIDRα1.5, CIDRα1.5, CIDRα1.4 | 4μg | 10 |
| | *Cocktail-semi-conserved-cVLPs* | cVLP-GA271, cVLP-GA273, cVLP-GA278, cVLP-GA282 | CIDRα1.8, CIDRα1.7, CIDRα1.4, CIDRα1.5 | | |
| | *Cocktail-diverse-cVLPs* | cVLP-GA268, cVLP-GA269, cVLP-GA272, cVLP-GA281 | CIDRα1.1, CIDRα1.6b, CIDRα1.7, CIDRα1.5a | | |
| **Mosaic-cVLPs** | *Mosaic-conserved-cVLPs* | cVLP-GA273-GA279-GA280-GA285-GA286 | CIDRα1.7, CIDRα1.4, CIDRα1.5, CIDRα1.5, CIDRα1.4 | 5μg | 10 |
| | *Mosaic-semi-conserved-cVLPs* | cVLP-GA271-GA273-GA278-GA282-GA284 | CIDRα1.8, CIDRα1.7, CIDRα1.4, CIDRα1.5, CIDRα1.6 | | |
| | *Mosaic-diverse-cVLPs* | cVLP-GA268-GA269-GA272-GA275-GA281 | CIDRα1.1, CIDRα1.6b, CIDRα1.7, CIDRα1.4, CIDRα1.5a | | 5 |
| **Recombinant cocktails** | *Recombinant cocktail-conserved* | GA273, GA279, GA280, GA285, GA286 | CIDRα1.7, CIDRα1.4, CIDRα1.5, CIDRα1.5, CIDRα1.4 | 5μg | 5 |
| | *Recombinant cocktail-semi-conserved* | GA271, GA273, GA278, GA282, GA284 | CIDRα1.8, CIDRα1.7, CIDRα1.4, CIDRα1.5, CIDRα1.6 | | |

|  |  |  |  |  |
| --- | --- | --- | --- | --- |
| Recombinant<br>*cocktail-diverse* | GA268, GA269,<br>GA272, GA275,<br>GA281 | CIDRα1.1, CIDRα1.6b,<br>CIDRα1.7, CIDRα1.4,<br>CIDRα1.5a | | |

First, we assessed IgG responses to the mosaic and cocktail cVLP vaccines (Fig 3A). IgG levels were highest (typically >7000 MFI) against variants included in the vaccines. Cross-reactive responses were mainly observed to CIDRα1 variants belonging to the same subgroup as used in the vaccine, reflecting a correlation with sequence similarity (Fig 3B). There were no major differences in the responses to mosaic and cocktail cVLP vaccines and no clear indication of induction of broadly reactive antibodies. Significant inhibition of EPCR binding was correlated with IgG reactivity levels (Fig 3C), and mostly but not always seen with reactivity >6000 MFI. Near complete neutralization was only observed for variants included in the vaccines.

**Fig 3.**
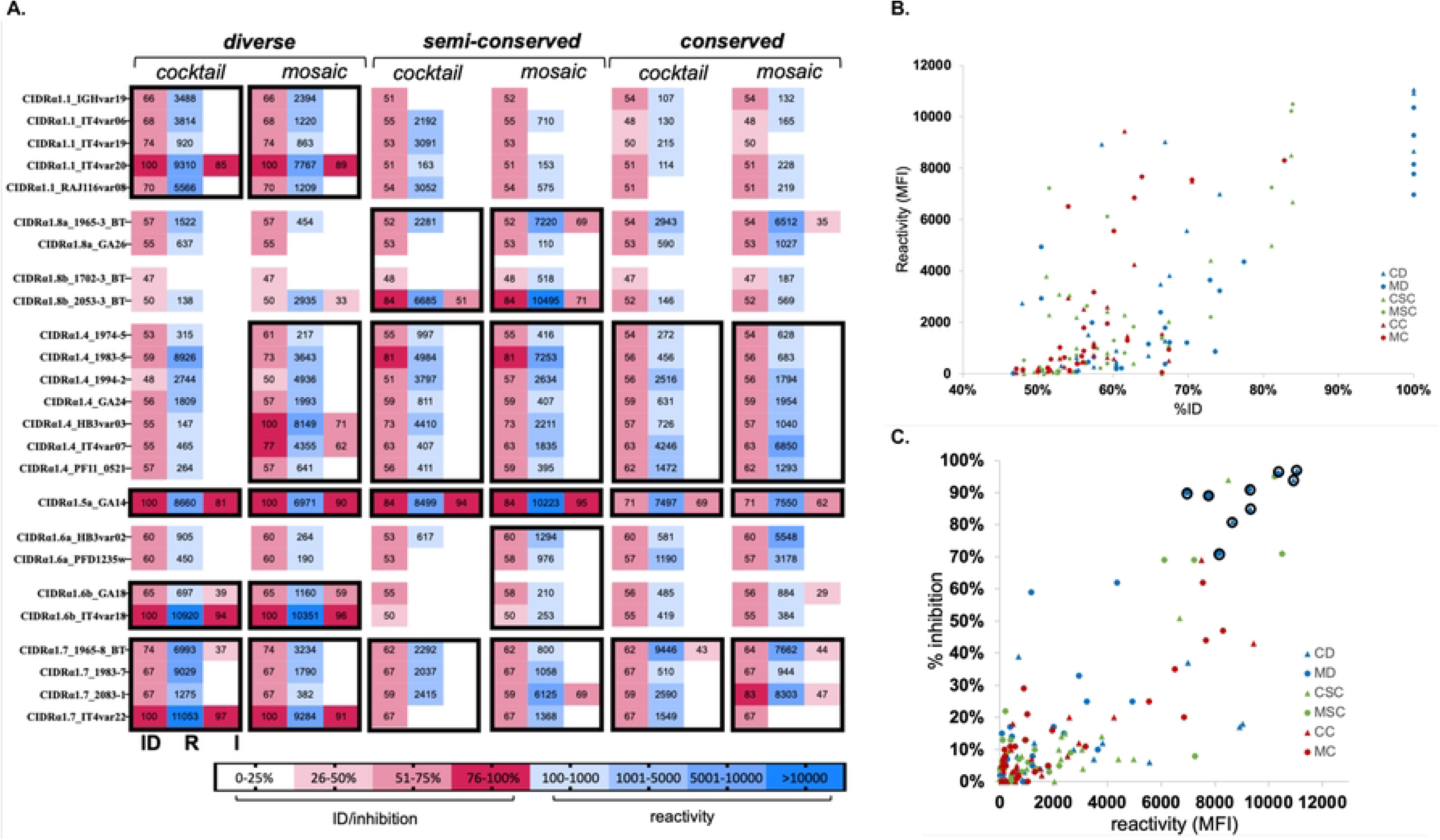
Reactivity and EPCR-binding inhibition of elicited antibodies. **A**. Heat map showing antibody responses to 25 CIDRα1 protein variants (rows). For each vaccination group (columns), percentage of identity (ID) between vaccine and test protein, IgG’s reactivity (R) towards the test protein variant measured as mean fluorescence intensity (MFI) and the IgG’s EPRC-binding inhibitory ability (I) are reported. For each test antigen variant, the ID score refers to the highest pairwise identity observed between the test antigen and any of the five CIDRα1 variants in the given vaccine. Black rectangles identify CIDRα1 subgroups (CIDRα1.1-8) to which variants in the vaccine belong. **B**. Correlation of sequence identity (%ID) and IgG reactivity (MFI). Each symbol represents reactivity to and percentage of identity with one of the 25 CIDRα1 test proteins. Spearman’s rank correlation = 0,6447, p-value <0.0001, alpha = 0,05. **C**. Correlation of reactivity (MFI) and EPCR-binding inhibition ability of pooled IgG from each vaccination group. Each symbol corresponds to inhibition and reactivity against one of 25 test CIDRα1 proteins. Spearman’s rank correlation = 0.6626, p-value <0.0001, alpha = 0,05. CIDRα1 test variants showing 100% sequence identity to vaccine variants are circled. Abbreviations: CD, cocktail-diverse; MD, mosaic-diverse: CSC, cocktail-semi-conserved; MSC, mosaic-semi-conserved; CC, cocktail-conserved; MC, mosaic-conserved.

Immune responses elicited by homotypic cVLPs (S5 Fig) mirrored the pattern observed for cocktail and mosaic cVLP vaccines, suggesting that the response observed in the latter is the result of cumulative dominant responses against each CIDRα1 variant present in the vaccine. In comparing the cVLP and recombinant protein cocktail vaccines, the reactivity and EPCR-binding inhibition patterns were similar, although the cocktail cVLP vaccines had weak but significant tendency to elicit higher and broader reactivity (S6 Fig), despite containing one less CIDRα1 variant.

## Discussion

Here we explored if immunization with AP205 cVLPs presenting a mosaic of different CIDRα1 sequence variants would facilitate the elicitation of broadly reactive antibody responses across the CIDRα1 protein family. The rationale was that the cVLP display of sequence-diverse CIDRα1 proteins sharing the same basic fold and a few surface-exposed residues placed at and around the EPCR binding site, would facilitate direct activation of B-cells bearing B-cell receptors binding to the antigens shared epitope. Conversely, activation of variant-specific B-cells would be less likely.

We hypothesized that recombinant CIDRα1 domains share potential B-cell epitopes as they, despite their extensive sequence diversity, bind EPCR through a similar structural complementarity and biochemical features, including a single conserved phenylalanine in the center of the EPCR binding site. As CIDRα1 sequences group into six major subgroups, we included CIDRα1 variants representative of five subgroups for the most diverse mosaic cVLP. To promote immune responses towards the EPCR binding site, we designed mosaics of five CIDRα1 domains sharing a different proportion of specific amino acids at the binding site. In all three cases, we found no evidence of significantly broader reactivity across CIDRα1 proteins elicited by the mosaic cVLPs compared to the vaccinations with the cocktails of homotypic cVLPs. While the spatial distribution of different CIDRα1 variants on the mosaic particles cannot be determined, mass spectrometry and immune responses indicate the successful coupling of all variants, most likely in a mosaic composition.

Several recent studies have reported that vaccination with mosaic antigen nanoparticles compared to homotypic nanoparticles improve breadth of neutralizing antibody responses against influenza and SARS-CoV-19 variants [17–23]. These studies have mainly relied on nanoparticles formed by antigen fused to self-assembling ferritin or proteins. One study, however, exploited AP205 SpyC-cVLPs to present mosaics of influenza hemagglutinin trimers and found no difference in breadth of immune responses elicited by mosaic and cocktails of homotypic cVLPs. Without direct comparison of antigen and study designs, it is at this stage impossible to assess if certain vaccine design strategies are more likely to generate broadly reactive antibody responses. Thus, in the present study it is possible that AP205 display of N-terminally SpyTag-fused CIDRα1 proteins confers suboptimal presentation of common inhibitory epitopes, or that these are few or of lower immunogenicity compared to other less broadly shared epitopes. Indeed, most cross-reactive responses were non-inhibitory. However, reactivity and inhibition were strongly and significantly correlated, suggesting that lack of broad reactivity may be related to excessive affinity maturation of variant-specific B-cells. This calls for an improved antigen design and display, taking into account heterogenicity of mosaic cVLPs, antigen spacing and orientation, thought to affect activation of B cells tolerating antigenic variability [18,24–26]. Including an increased number of variants or revising immunization dosing and timing, may be additionally important for broadly reactive antibodies to develop [27,28]. Advanced artificial intelligence-aided epitope or antigen design and a better insight to the molecular mechanisms and characteristics of naturally acquired broadly reactive and neutralizing antibodies may inform new vaccine conceptualization and development.

As previously reported, we observed a slightly higher reactivity of antibodies elicited by immunizations with cVLP-displayed CIDRα1 proteins, compared to immunizations with the corresponding recombinant CIDRα1 proteins [6]. Also in line with other studies, cross-reactivity was elicited by all our vaccine designs, although it was restricted to CIDRα1 variants within the same subgroup included in the vaccines [6–8]. This suggests that a polyvalent vaccine eliciting cross-reactive responses within each of the six main CIDRα1 subgroups is possible, although current data makes it is difficult to assess the CIDRα1 antigen complexity required to gain this desired response.

## Materials and methods

### Design, expression and purification of recombinant CIDRα1

CIDRα1 sequences were designed with domain borders corresponding to amino acid 499-719 of the HB3VAR03 [5]. SpyCatcher (SpyC) (genebank: OK422508.1) and Strep-tag II proprietary peptide sequences were genetically fused through a flexible linker (GSGS) to the N-and C-terminus of the selected CIDRα1 sequences, respectively. DNA sequences were synthesized by GeneArt and used for protein expression in *Spodoptera frugiperda* (Sf9) cells, as previously [6]. In brief, High Five™ cells were infected with recombinant virus. Soluble recombinant CIDRα1 proteins were affinity purified in StrepTrapTM HP/XT 1 mL columns (Cytiva), and dialyzed in 1X PBS pH 8.0. Purity, absence of aggregates and correct molecular weight of the purified products were confirmed in SDS-PAGE prior coupling to cVLPs. EPCR binding ability of recombinant CIDRα1 was validated in ELISA, as previously described [6].

### Formulation and quality assessment of CIDRα1-cVLP vaccines

AP205 cVLPs presenting one SpyTag (SpyT) (genebank: OK545878.1) per capsid unit were prepared as previously described [6,15,16]. According to the different combinations (Fig 2A), recombinant CIDRα1 proteins and assembled SpyT-cVLPs were mixed in a 1:1 molar ratio, incubated for 2 hours at RT and dialyzed overnight using 1000 kDa MWCO dialysis membrane (SpectraPor) in 1X PBS, 200 mM sucrose, 15 mM Tris (pH 8.0). A second dialysis at 4°C in fresh buffer was carried out for 4 hours to ensure removal of non-coupled proteins. Post-dialysis samples were subjected to a spin test to assess stability, as previously indicated [16]. Pre/post-dialysis and post-spin test samples were loaded on SDS-PAGE, under reducing conditions (Fig 2B).

Densitometric analysis of the SDS-PAGE allowed estimation of SpyC-CIDRα1 coupling efficiency of homotypic vaccines, defined as percentage of coupled cVLP subunits over total cVLP subunits, in ImageLab [15,16].

Successful coupling of recombinant CIDRα1 for mosaic cVLPs was assessed in a bottom-up, label-free, MS approach. Samples were analyzed by the Proteomics Core facility of the Technical University of Denmark (DTU), following standard protocol used for bottom-up and label free proteomics analysis (*supplementary* methods).

DLS analysis (DynaPro Nanostar, Wyatt technology) of cocktail-and mosaic-cVLP formulations was performed at room temperature, following sample spinning as described above. 20 acquisitions of 5 seconds were run for each cVLP preparation. Estimated particle diameter and percentage of polydispersity (%Pd) were computed using Wyatt DYNAMICS software, as previously described [16].

### Mice immunizations

Homotypic, cocktails-and mosaic-cVLP vaccines were formulated without adjuvant in 1X PBS, 200 mM sucrose, 15 mM TRIS, pH 8.0. Each mouse vaccinated with homotypic cVLPs received 2 μg of coupled CIDRα1 per immunization. A dose of 1 μg of each recombinant CIDRα1 protein was administered per mouse per immunization, in the case of cocktails and mosaics formulations. For mosaic cVLPs the total amount of coupled CIDRα1 injected was 5 μg, while in the case of cocktail cVLPs it was 4 μg, due to precipitation or sample loss of one homotypic cVLPs sample in each of the three mixes. In total, 85 female BALB/c mice, 9 weeks old were distributed in 12 different vaccination groups (Table 1) and immunized intramuscularly twice, three weeks apart. Serum was collected two weeks following booster dose.

Cocktails of soluble proteins were obtained by mixing 1 μg of each individual CIDRα1 (5 μg in total), according to the composition of conserved, semi-conserved and diverse cVLP vaccines. Formulations were diluted in 1X PBS and administered with Addavax^TM^ (Invivogen) (1:1). A total of 15 mice (Table 1) were immunized intramuscularly and serum was collected as described above.

### IgG reactivity and EPCR binding inhibition

Total IgG was purified in Poly-Prep Chromatography columns (Bio-Rad Laboratories, Inc) packed with Gammabind Plus Sepharose (Cytiva), according to manufacturer’s protocol. Purified IgGs were pooled according to vaccination group and diluted in ABE buffer (1X PBS, 0.02% Tween20, 0.1% BSA) to a final concentration of 0.25 mg/ml. Reactivity and EPCR-binding inhibition was assessed towards a custom-made panel of recombinantly produced CIDRα domains conjugated to Luminex microspheres, using Bio-Plex™ 200 system (Bio-Rad), as previously reported [6,29]. Briefly, 50 μL/well of plex beads were distributed in a MultiScreenHTS BV Filter Plate (1.2 μm Hydrophilic Low Protein Binding Durapore Membrane, Merck) and washed three times with 120 μL of ABE buffer. 50 μL of sample diluted IgGs were added to the wells and after incubation in the dark on a shaking platform (30 s at 600 rpm followed by 30 min at 300 rpm), the plate was washed as before. IgG reactivity was detected by adding 50 μL/well R-Phycoerythrin-(PE)-conjugated AffiniPure F(ab’)_2_ Fragment Goat Anti-Mouse antibody (Jackson ImmunoResearch, Code Number: 115-116-146, 1:3500) following incubation and washes, as before. Prior to acquisition, 50 μL/well of ABE buffer was aliquoted and the plate was shaken one last time at 300 rpm for 1 minute. For EPCR-binding inhibition assessment an additional incubation with 50 μL of biotinylated EPCR to a final concentration of 4 μg/mL was performed, prior to incubation with PE-conjugated Streptavidin-R (SIGMA-ALDRICH). Computation of reactivity and inhibition values is reported in the *supplementary methods*.

### Data analysis

Statistical analyses were conducted exploiting specific built-in functions of GraphPad Prism software (Dotmatics) in all cases. Specific analyses are reported in the corresponding *Results* section.

## Acknowledgments

The authors wish to express their gratitude to Akiko Shiraishi for technical support in production of recombinant CIDRα1 proteins, Lærke Lillelund for the assistance with IgG purification, Sai Raghavan for his insights and advice, Louise Goksøyr, Anna Kathrine Okholm and Anne Corfitz for mice immunizations. We thank Lundbeck Foundation (R344-2020-934), the Independent research Fund Denmark (9039-00285A) and Kirsten of Freddy Johansens Fond for financial support.

## Supporting information

**S1 Fig.**
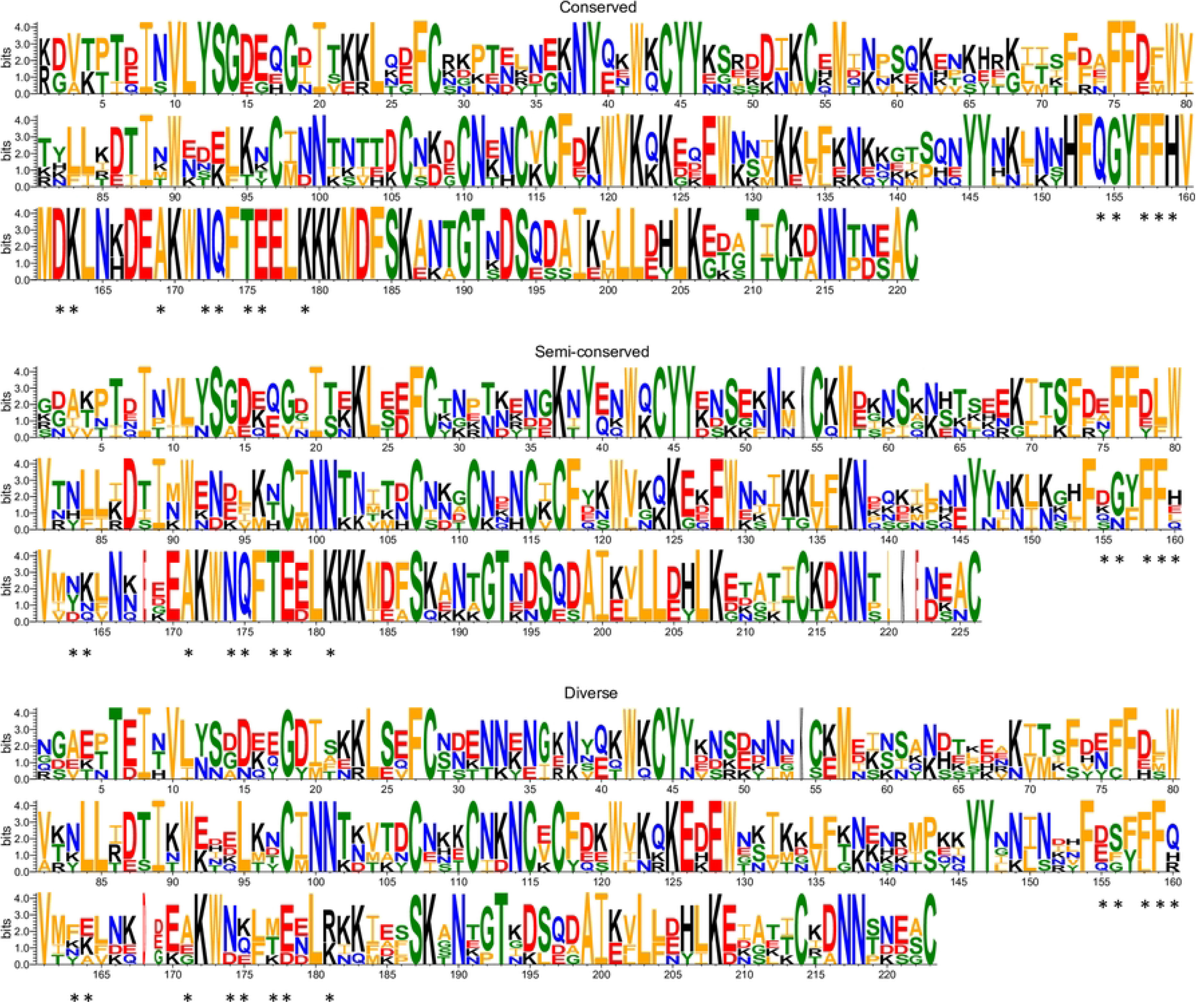
Sequence conservation LOGO of CIDRα1 variants included in the vaccines. Logo were generated in WebLogo3 by alignment of the five CIDRα1 selected according to the respective vaccination strategy (*conserved*, *semi-conserved* and *diverse*). Asterisks indicate the 13 specified ammino acid positions constituting the putative epitope.

**S2 Fig.**
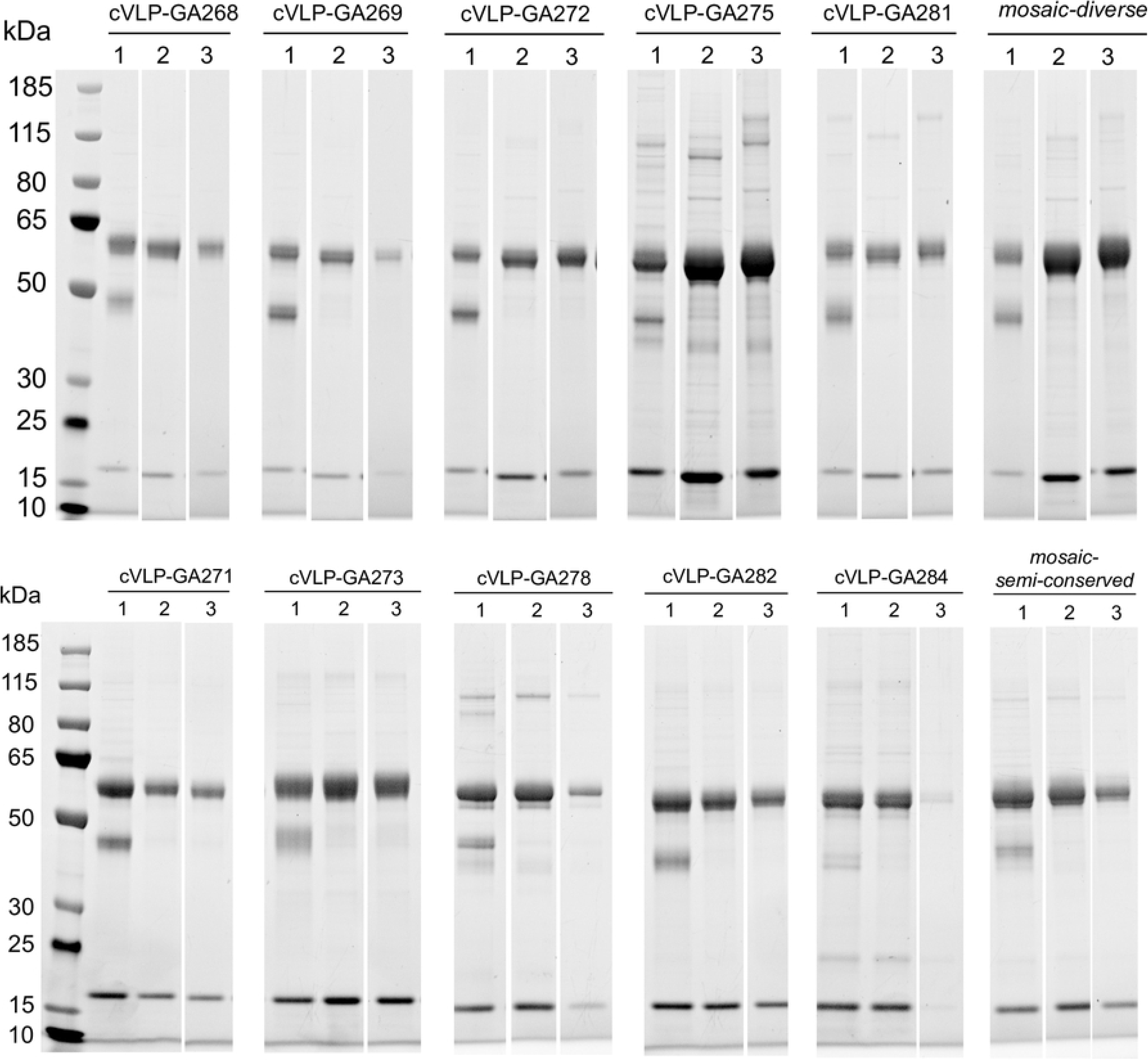
SDS-PAGE of 10 homotypic cVLPs, *mosaic-diverse*-cVLPs and *mosaic-semi-conserved*-cVLPs. First lane (1) represents the samples pre-dialysis. The bands correspond to Tag-cVLP (16.5 kDa), unbound Catcher-CIDRα1 (∼40 kDa) and coupled CIDRα1-cVLP (∼56 kDa). The second lane (2) represents the samples post-dialysis and before spin test. The third lane (3) represents the samples post-dialysis and post-spin test. cVLP-GA275 and cVLP-GA284 were excluded from the study. The amount of cVLP-GA275 sample recovered after dialysis was insufficient to proceed with further analyses and immunization, while cVLP-GA284 demonstrated propensity to aggregation and instability upon performance of the spin test.

**S3 Fig.**
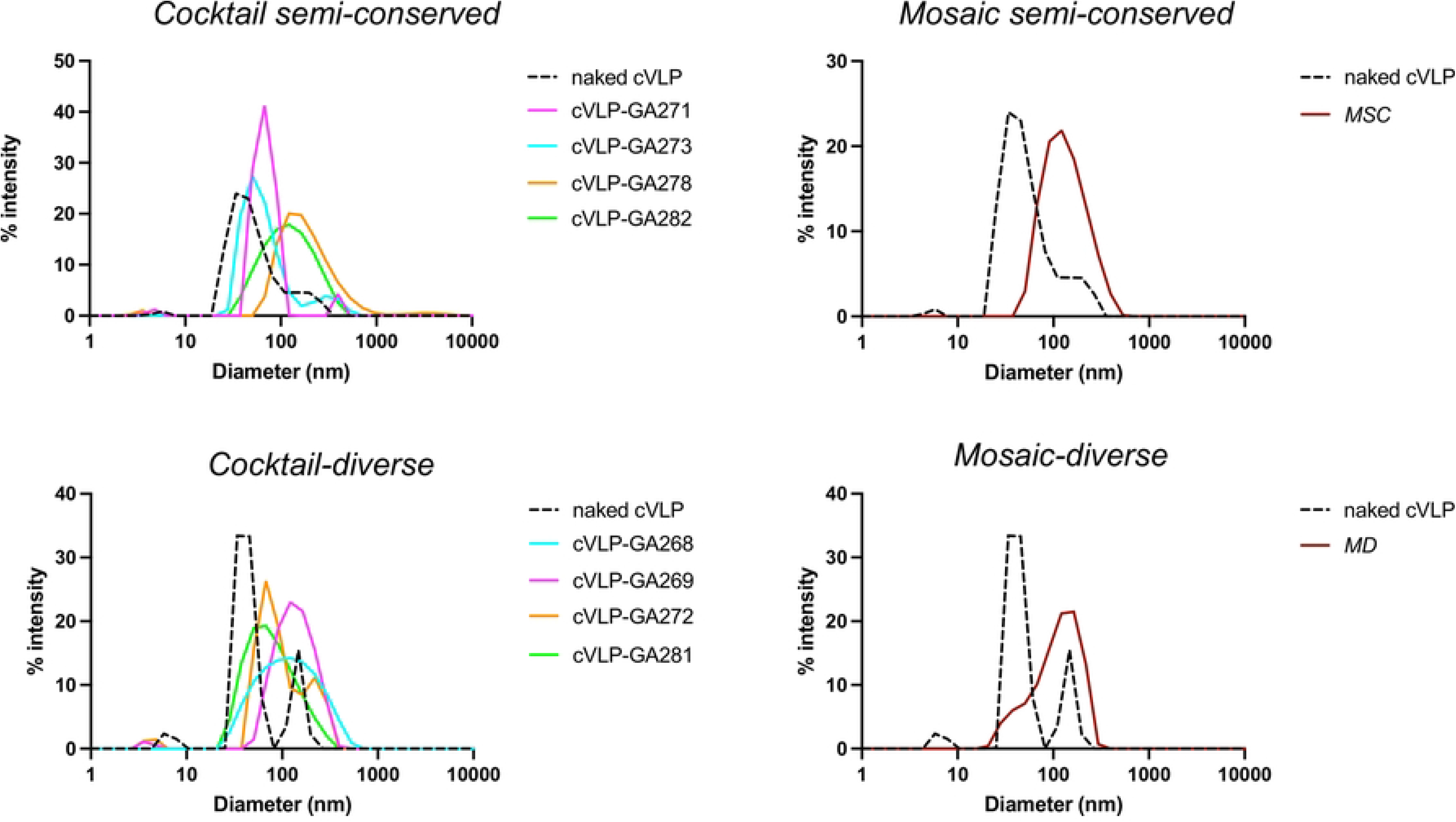
Dynamic-Light-Scattering (DLS) analysis of *cocktail-semi-conserved/diverse*-cVLPs and *mosaic-semi-conserved/diverse-*cVLPs. Naked SpyT-cVLPs (dashed line) predominant population shows 65.7 nm with 81.2% Pd in the *cocktail*-and -*mosaic*-*semi-conserved*-cVLPs and 41.9 nm with 20.5% Pd in the *cocktail-and* -*mosaic*-*diverse*-cVLPs. *Cocktail-semi-conserved-*cVLPs indicate a population size of 63.3-138.2 nm with 22.2-58.8% Pd, *mosaic-semi-conserved-*cVLPs show 146.8 nm and 53.1% Pd, *cocktail-diverse-*cVLPs 86.6-147.4 nm with 39.4-70.4% Pd and *mosaic-diverse-*cVLPs 120.3 nm with 49% Pd. Abbreviations: MD, mosaic-diverse; MSC, mosaic-semi-conserved.

**S4 Fig.**
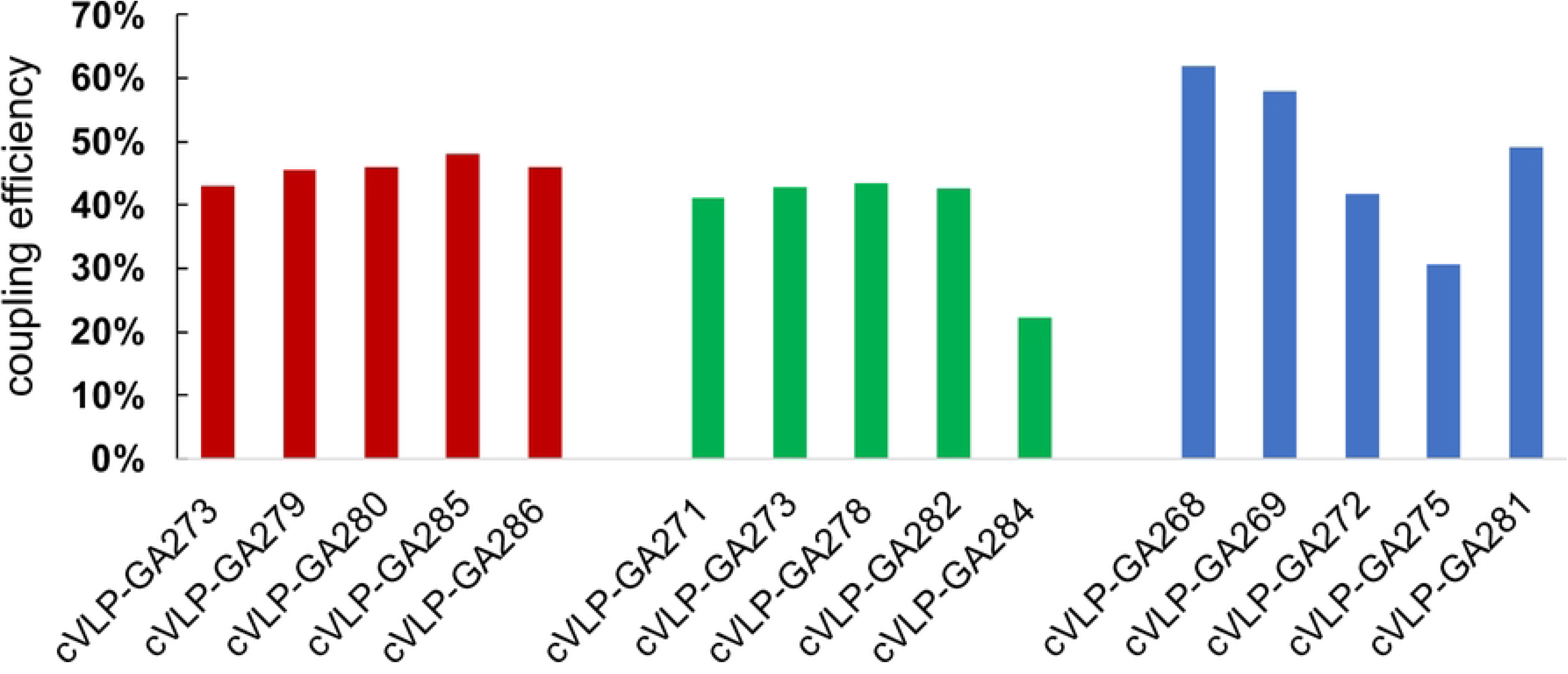
Coupling efficiency estimated by densitometry analysis of homotypic cVLPs.

**S5 Fig.**
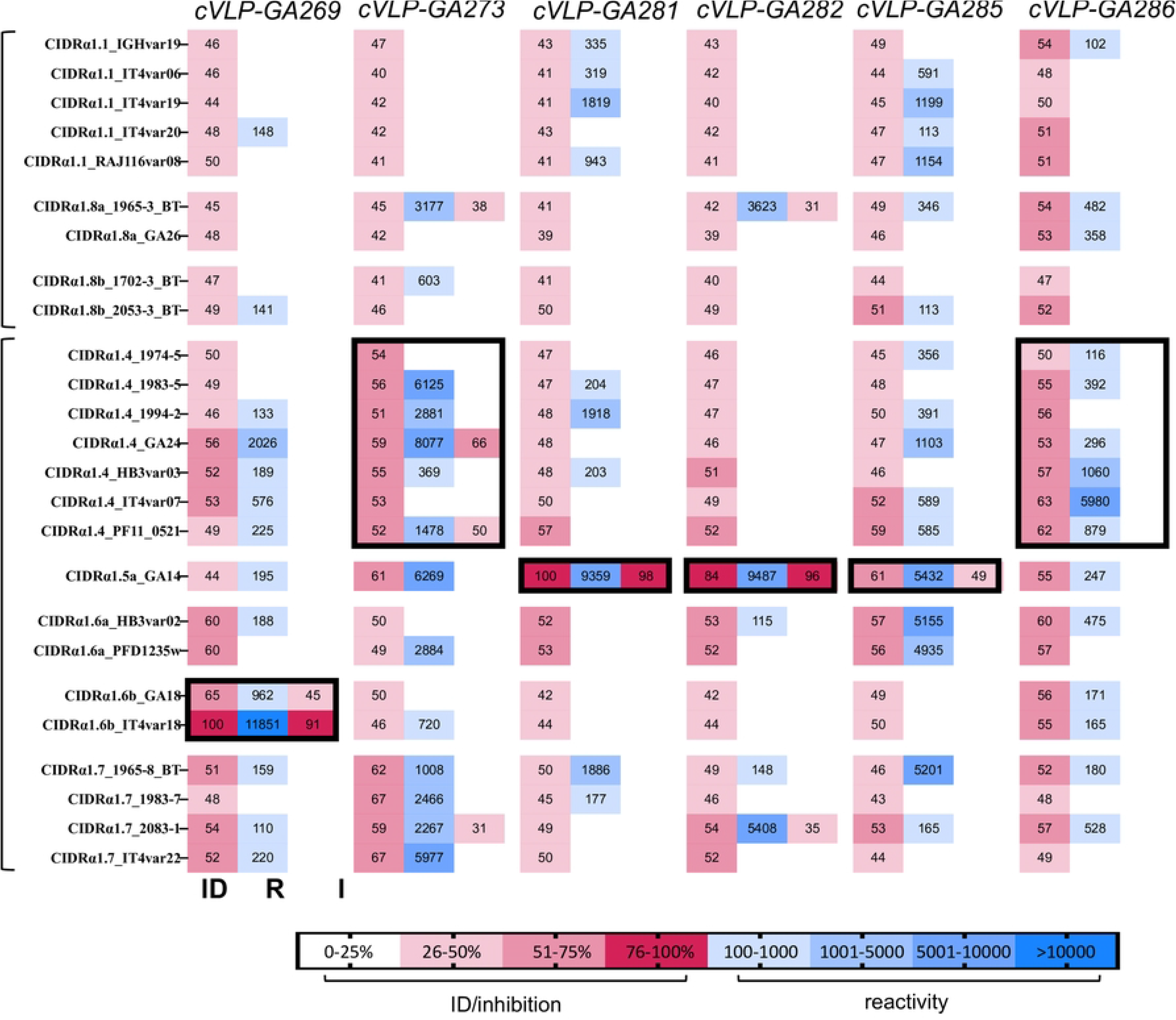
Reactivity and EPCR-binding inhibition of antibodies elicited with homotypic cVLPs vaccines. Heat map showing antibody responses to 25 CIDRα1 protein variants (rows). For each vaccination group (columns), percentage of identity (ID) between vaccine and test protein, IgG’s reactivity (R) towards the test protein variant measured as mean fluorescence intensity (MFI) and the IgG’s EPRC-binding inhibitory ability (I) are reported. Black rectangles identify CIDRα1 subgroups (CIDRα1.1-8) to which variants in the vaccine belong.

**S6 Fig.**
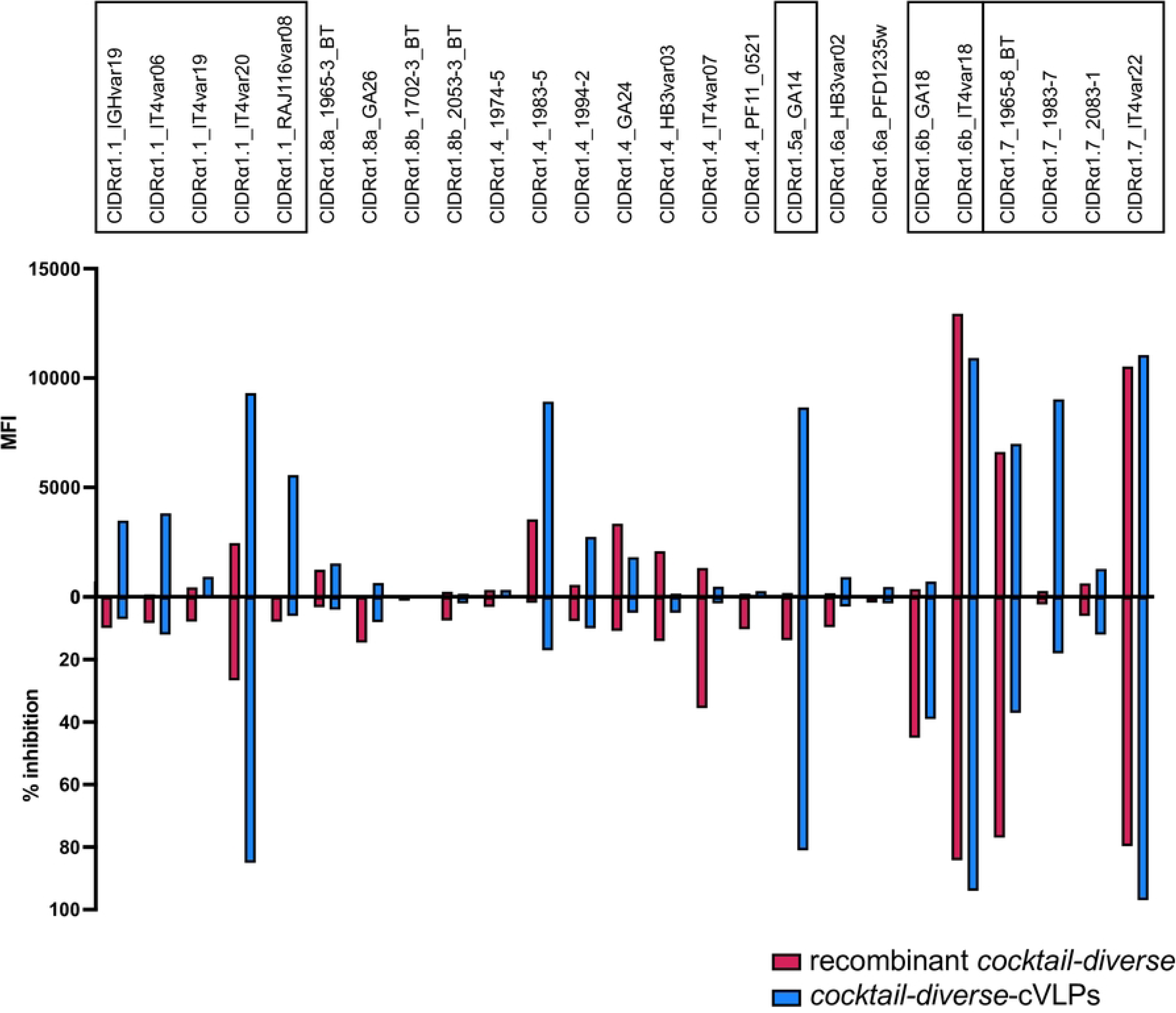
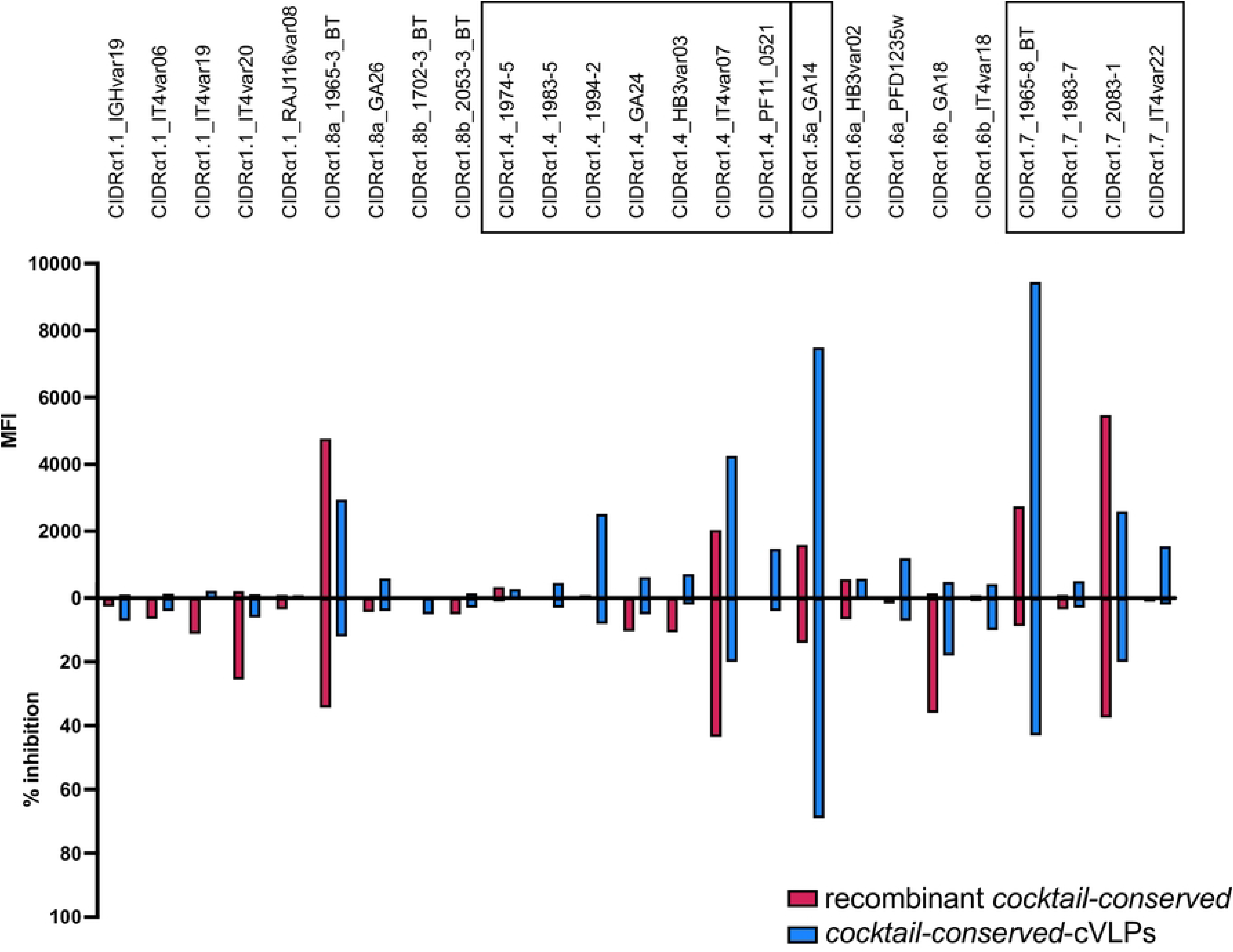
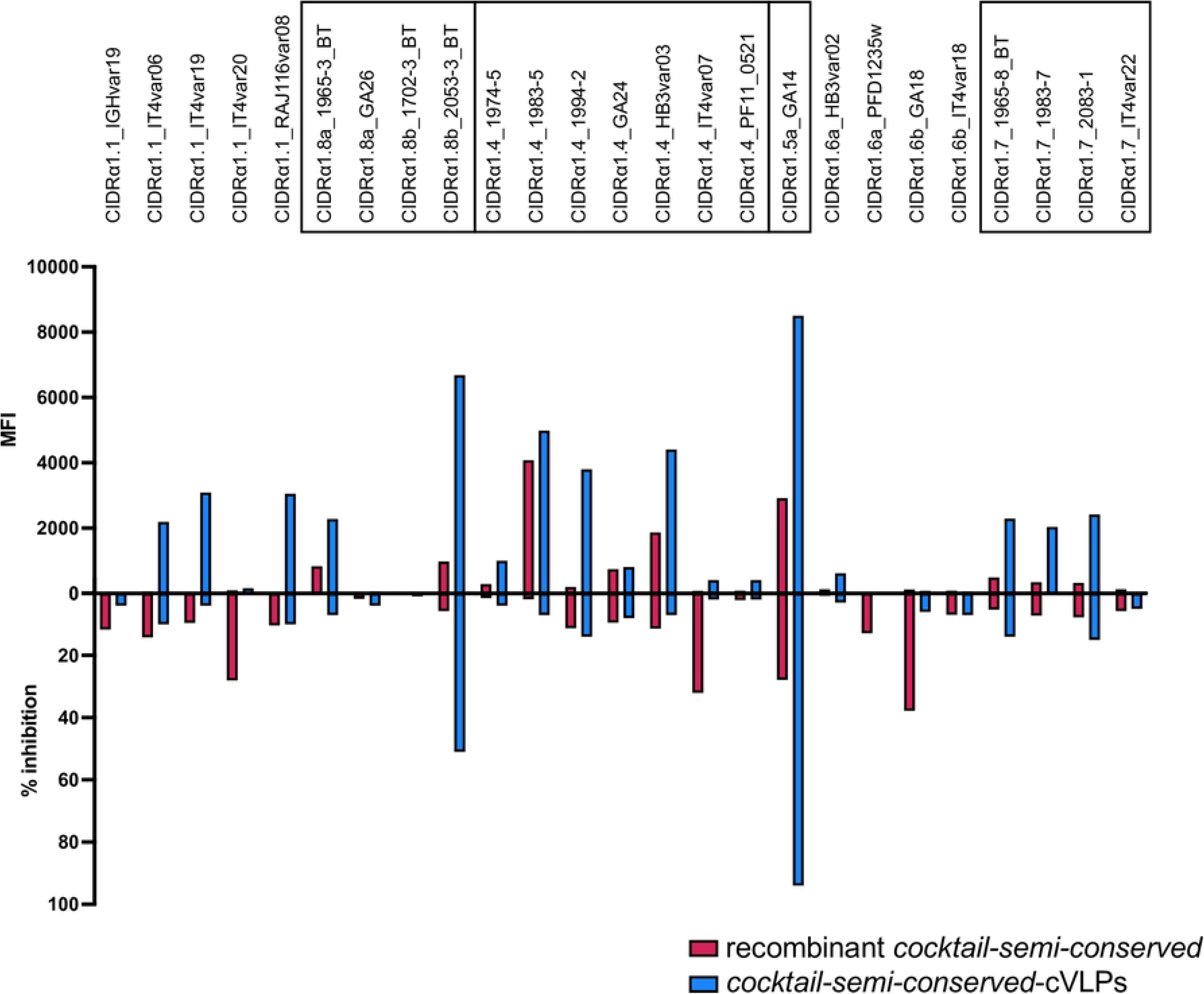
Reactivity and EPCR-binding inhibition of antibodies elicited by recombinant cocktails. Antibody responses were tested against the same 25 CIDRα1 protein variants panel used to test immunogenicity of cVLP-based vaccines. Data analysis was carried out as before. Black rectangles identify CIDRα1 subgroups (CIDRα1.1-8) to which variants in the vaccine belong. Wilcoxon matched-pairs signed rank test resulted in significant reactivity differences in all three cases (recombinant vs *cocktail*-*diverse-*cVLPs: p-value = 0,0088; recombinant vs *cocktail*-*semi-conserved-*cVLPs: p-value <0,0001; recombinant vs *cocktail*-*conserved-*cVLPs: p-value = 0,0025). No significant differences in inhibition were found exploiting the same statistical test.

**S1 file. Supplementary methods.**

**S2 file. Recombinant protein sequences.**

